# Infection of *Borrelia burgdorferi* sensu lato of small mammals in Yunnan Province, China

**DOI:** 10.1101/714980

**Authors:** Zhi-hai He, Bao-gui Jiang, Zi-hou Gao, Zong-ti Shao, Yun Zhang, Zheng-xiang Liu, Yu-qiong Li, En-nian Pu, Li Tang, Ming-guo Yao, Na Jia, Michael E. von Fricken, Jia-fu Jiang, Wu-chun Cao, Chun-hong Du

## Abstract

**Background:** Lyme disease is caused by *Borrelia burgdorferis*ensulato (BBSL) which is usually found in wild and domestic mammals worldwide. Human cases of *B. burgdorferi* infections have been identified in China, but little direct surveillance of potential rodent reservoirs has been performed in Yunnan Province, Southwestern China. Yunnan Province is a tropical area with a diverse topographic range and sustains a high biodiversity of small mammals that could potentially play an important role in the transmission of a variety of *B. burgdorferi*genospecies.

**Methods:** 3659 small mammals were captured in 159 sample siteslocated 23 countries inYunnan Province and screened for BBSL infection by nested PCR based on 5S-23S rRNA intergenic spacer gene of BBSL.Univariate and multivariate forward stepwise logistic regression analysis was used to access the association between infections and related risk factors.

**Results:** Infection with BBSL was confirmed in 3.99%(146/3659) of small mammals. Significant differences in prevalence rates of BBSL were observed at varying landscape types and altitudes.Small mammals in forested areas had higher prevalence rates than other landscape types as did small mammals found at altitudes greater than 2500 meters. The 5S-23S rRNA intergenic spacergene revealed that there were 5 genotype of BBSL, including *B. afzelii, B. burgdorferisensustricto, B.japonica, B.gariniiand B.valaisiana*, which demonstrate the genetic diversity and regional distribution.

**Conclusions:** There exists a wide distribution and genetic diversity of endemic BBSL in Southwestern China, warranting further investigations and monitoring of clinical disease in individuals presenting with symptoms of Lyme disease in these areas.

**Author summary:** Lyme disease is caused by *Borrelia burgdorferi* sensu lato (BBSL) which is usually found in wild and domestic mammals worldwide. Human cases of *Borrelia burgdorferi* sensu lato infections have been identified in China, but little direct surveillance of potential rodent reservoirs has been performed in Southwestern China. This study documents potential small mammal reservoir hosts collected from a large of sample sites from different landscape types and altitudes, with PCR and sequencing identifying the wide distribution and genetic diversity of endemic *Borrelia burgdorferi* sensu lato in Southwestern China. This was the first report that *B. japonica* was detected in *Apodemus draco* and *Niviventer excelsior* in China. This study adds to body of literature on *Borrelia burgdorferi* sensu lato in China. This work will provide insight regarding small mammals to target for surveillance and we access the association between gender, developmental stage of rodents, environmental landscape and altitude to better prevent human exposure.

## Introduction

Lyme borreliosis (LB) is the most commonly reported vector-borne disease across Europe, North America and Asia[1–4]. The causative agents of LB fall within the species complex *B. burgdorferi* sensu lato (BBSL), and is responsible for a wide spectrum of clinical symptoms. Anti-Borrelia antibodies in rats and humans have been reported in 9 counties and 4 counties of Yunnan, respectively. While there have been documented reports of human cases of Lyme disease in southwestern China[4], the only information on the prevalence of BBSL in rodent reservoirs came from one study, where a majority of rodents were trapped indoors[5]. Yunnan Province is of particular interest given its wide topographic range and high level of small mammal biodiversity, many of which may potential reservoirs for BBSL. We performed a systematic field investigation on the prevalence of BBSL infections in a large quantity of rodents sampled in 23 countries in Yunnan Province, and then analyzed the distribution and genetic diversity of BBSL, as well as the association between infections and suspected risk factors. This study aims to evaluate the role that small mammals play in the transmission of BBSL across Yunnan Province.

## Materials and methods

### Ethics statement

The research protocol for trapping wild small animals and collecting samples was approved by the Animal Subjects Research Review Boards at the Yunnan Institute of Endemic Diseases Control and Prevention (2013-003), in accordance with the medical research regulations of China and the Regulation of the People’s Republic of China for the Implementation of the Protection of Terrestrial Wildlife.

### Collection of small mammal samples

From 2011 to 2016, small mammals were captured using animal snap traps set at agricultural, forested, and residential areas at 159 sample sites from 23 counties ranging from 530 to 4300m in Yunnan Province (Table 1). Two hundred snap traps per sample site were placed for three consecutive nights and checked daily. Mammal species were identified according to external morphology, fur color, measurements and visible characters of dentition. Each animal’s sex, developmental stage, and location was recorded at the time of sample processing. After identification of species, spleen tissues were removed from the animals and stored in liquid nitrogen until tested. For unidentified species in the field, the craniums were brought to the laboratory for further identification.

**Table 1.** Prevalence of BBSL in small mammals from different survey sites.

| Counties | Sampling site | No. of tested | No. of positive for BBSL (%) | B.a | B.b | B.g | B.j | B.v |
| --- | --- | --- | --- | --- | --- | --- | --- | --- |
| Gongshan | S1 | 711 | 61 (8.58) | 39 | 22 | 0 | 0 | 0 |
| Deqin | S2 | 662 | 52 (7.85) | 20 | 1 | 30 | 0 | 1 |
| Shangri-la | S3 | 106 | 0 (0) | 0 | 0 | 0 | 0 | 0 |
| Fugong | S4 | 134 | 1 (0.75) | 0 | 1 | 0 | 0 | 0 |
| Weixi | S5 | 239 | 8 (3.35) | 6 | 2 | 0 | 0 | 0 |
| Yulong | S6 | 224 | 1 (0.45) | 1 | 0 | 0 | 0 | 0 |
| Jianchuan | S7 | 112 | 0 (0) | 0 | 0 | 0 | 0 | 0 |
| Lushui | S8 | 114 | 0 (0) | 0 | 0 | 0 | 0 | 0 |
| Yunlong | S9 | 177 | 2 (1.13) | 0 | 0 | 0 | 2 | 0 |
| Tengchong | S10 | 39 | 1 (2.56) | 0 | 0 | 0 | 0 | 1 |
| Yingjiang | S11 | 38 | 0 (0) | 0 | 0 | 0 | 0 | 0 |
| Yongde | S12 | 215 | 3 (1.40) | 2 | 0 | 0 | 0 | 1 |
| Yunxian | S13 | 68 | 1 (1.47) | 0 | 0 | 0 | 0 | 1 |
| Jinggu | S14 | 76 | 1 (1.32) | 1 | 0 | 0 | 0 | 0 |
| Ninger | S15 | 88 | 0 (0) | 0 | 0 | 0 | 0 | 0 |
| Yiliang | S16 | 94 | 6 (6.38) | 6 | 0 | 0 | 0 | 0 |
| Shiping | S17 | 128 | 6 (4.69) | 5 | 1 | 0 | 0 | 0 |
| Mile | S18 | 110 | 2 (1.82) | 2 | 0 | 0 | 0 | 0 |
| Mengzi | S19 | 22 | 0 (0) | 0 | 0 | 0 | 0 | 0 |
| Jinping | S20 | 27 | 0 (0) | 0 | 0 | 0 | 0 | 0 |
| Wenshan | S21 | 17 | 0 (0) | 0 | 0 | 0 | 0 | 0 |
| Menghai | S22 | 177 | 1 (0.56) | 1 | 0 | 0 | 0 | 0 |
| Mengla | S23 | 81 | 0 (0) | 0 | 0 | 0 | 0 | 0 |
| Total |  | 3659 | 146 (3.99) | 83 | 27 | 30 | 2 | 4 |

### DNA extraction and PCR analysis

DNA was extracted from spleen tissue using the DNA blood and tissue kit (Tiangen Biotechnique, Beijing, China) according to the manufacturer’s instruction. A nested PCR for the 5S-23S rRNA intergenic spacer gene of BBSL was done as previously described[6]. The PCR-positive amplicons were directly sequenced with an automated DNA sequencer (ABI PRISM 373; Perkin-Elmer, Norwalk, CT). Sequence analysis was carried out using a FASTA search on the Genbank database, with phylogenetic trees constructed using MEGA software, version 6.06[7]. The 5S-23S rRNA intergenic spacer gene of BBSL obtained in this study were deposited in Genbank under accession numbers MK333406-MK33427 and KP677523.1 respectively.

### Statistical analysis

Univariate analysis was used to access the association between gender, developmental stage of rodents, environmental landscape, altitude, and testing positive for BBSL using a chi-square test. All variables with a *P*-value of <0.05 from univariate analysis were entered into a multivariate forward stepwise logistic regression analysis. All analyses were conducted using SPSS (version 17.0, SPSS Inc. Chicago, IL).

## Results

A total of 3659 small mammals belonging to 57 species, 29 genera and 10 families from 5 orders were collected (Table 1). The *Apodemus draco* was the most common species (15.82%, 579/3659), followed by *Rattus tanezumi* (15.66%, 573/3659). A total of 146 (3.99%) rodentstested positive for BBSL, with *Ochotona gloveri* (33.33%, 1/3), *Sorex cylindricauda* (14.28%, 7/49), *Soriculusleucops* (14.94%,13/87), and *Rattus tuekkestanicus* (14.28%, 1/7) actively infected with BBSL (Table 2). The positive mammals originated from 14 out of 23 sample counties, including Deqin, Weixi, Yulong, Gongshan, Fugong, Jinggu, Tengchong, Yongde, Menghai, Yunxian, Shiping, Mile, Yiliang and Yunlong (Table 1), with Gongshan (S1) having the highest prevalence (8.58%), followed by Deqin (S2, 7.85%), and Yiliang (S16, 6.38%). The prevalence of BBSL in small mammals in forested landscapes, agricultural landscapesand residential landscapes were 5.19%, 3.14% and 0.63%, respectively. There was significant difference in prevalence of BBSL in small mammals at the altitude classes of <1500 meters, 1500-2500 meters, >2500 meters with 0.80%, 2.92% and 5.86%, respectively (χ^2^=43.089, *p*=0.001), and between different landscapes (χ^2^=14.945, *p*=0.001) as depicted in Table 3. The multivariate logistic regression analysis also revealed that samples found at altitudes greater than 1500 meters and in agricultural landscapes were more likely to be infected with BBSL (Table 4).

**Table 2.**
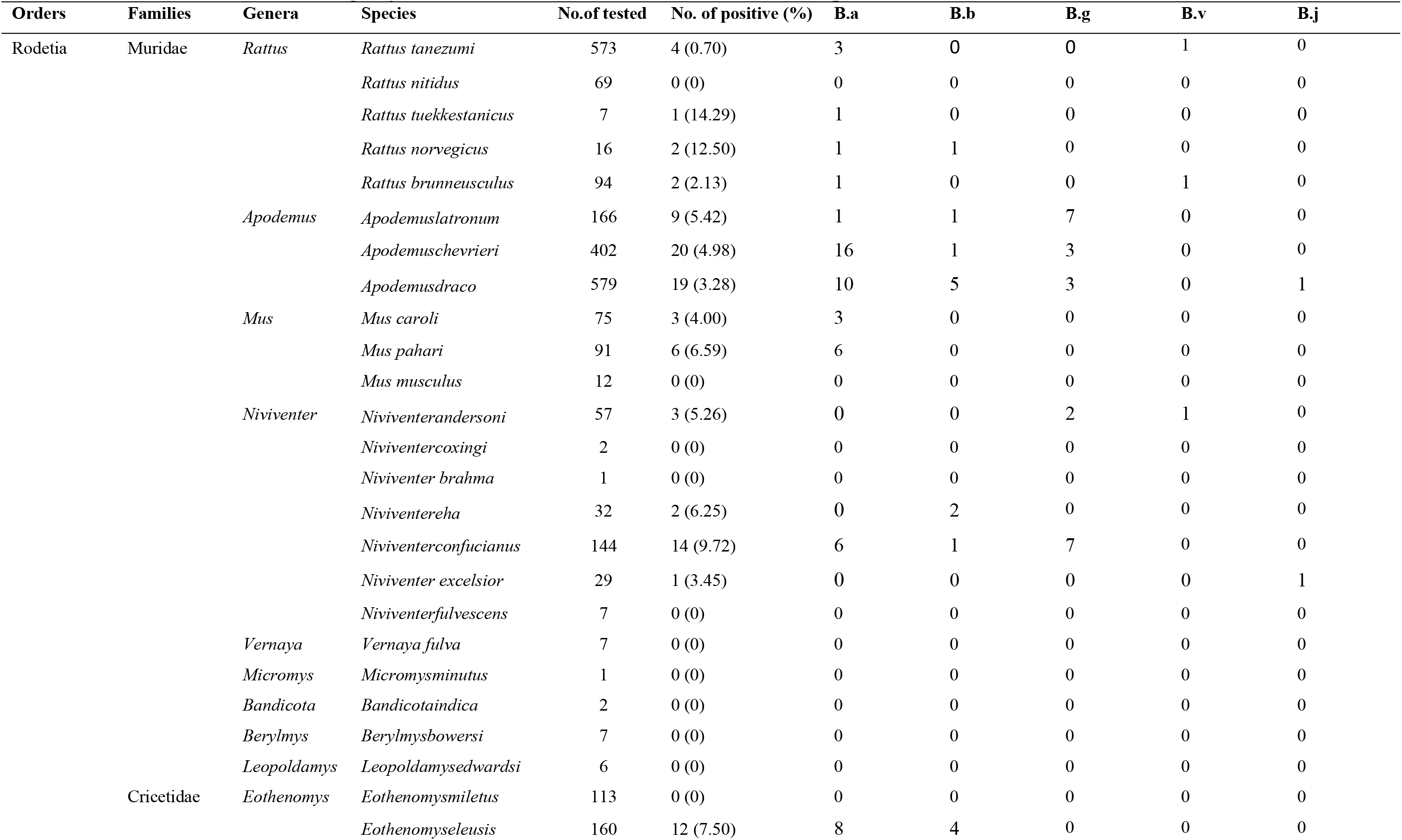
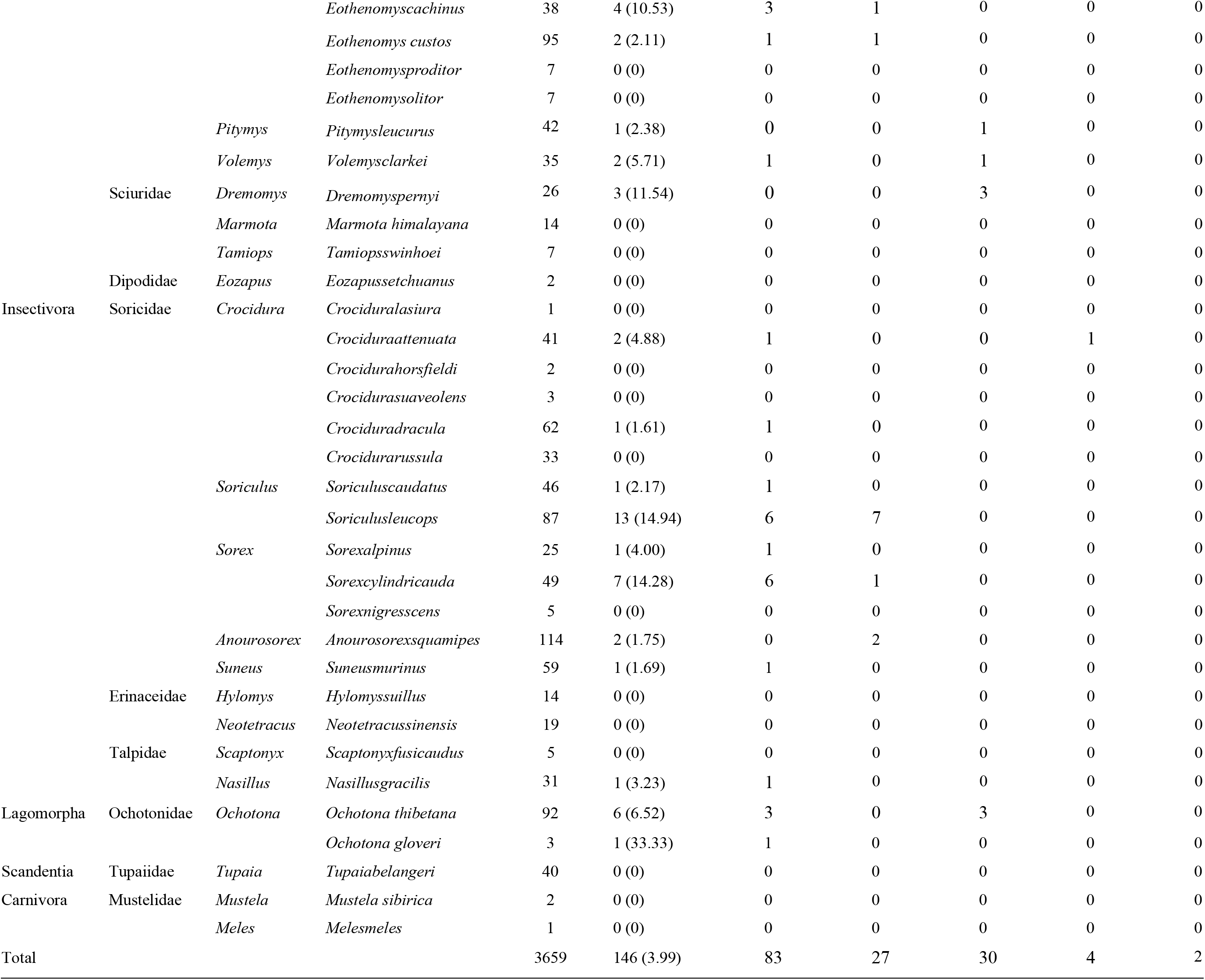
Prevalence of *Borrelia burgdorferi* sensu lato in small mammals of different species.

**Table 3.** Risk factors related to *Borrelia burgdorferi* sensu lato based on univariate analyses.

| variable | simple size | BBSL Infection |  |  |
| --- | --- | --- | --- | --- |
| | constituent ratio (%) | positive rate (%) | $\chi^2$ | P |
| altitude (m) |  |  |  |  |
| ~1500 | 868/3659 (23.72%) | 7/868 (0.81%) | 43.089 | 0.001 |
| 1500~2500 | 823/3659 (22.49%) | 24/823 (2.92%) |  |  |
| 2500~ | 1968/3659 (53.79%) | 115/1968 (5.84%) |  |  |
| gender |  |  |  |  |
| male | 1753/3659 (47.91%) | 81/1753 (4.62%) | 3.492 | 0.062 |
| female | 1906/3659 (52.09%) | 65/1906 (3.41%) |  |  |
| age |  |  |  |  |
| adult | 3337/3659 (91.20%) | 133/3337 (3.99%) | 0.002 | 0.964 |
| pubertal | 322/3659 (8.80%) | 13/322 (4.04%) |  |  |
| landscape |  |  |  |  |
| residential | 158/3659 (4.32%) | 1/158 (0.63%) | 14.945 | 0.001 |
| agricultural | 1786/3659 (48.81%) | 56/1786 (3.14%) |  |  |
| forest | 1715/3659 (46.87%) | 89/1715 (5.19%) |  |  |
| Total |  | 146/3659 (3.99%) |  |  |

**Table 4.** Risk factors related to BBSL based on multivariate logistic regression.

| Variable | OR (95% CI) | p |
| --- | --- | --- |
| **Altitude (m)** |  |  |
| <1500 | 1 |  |
| 1500-2500 | 4.524 (1.979~10.339) | <0.01 |
| >2500 | 24.489 (11.351~52.833) | <0.01 |
| **Landscape category** |  |  |
| **residential** | 1 |  |
| forest | 5.617 (0.769~41.052) | 0.089 |
| agricultural | 8.412 (1.150~61.528) | 0.036 |
| **forest** | 1 |  |
| residential | 0.178 (0.024~1.301) | 0.089 |
| agricultural | 1.497 (1.153~1.944) | 0.002 |

Sequencing was successful for all 146 positives amplicons samples. The comparative analysis with the BLAST program revealed that 83 samples were *B. afzelii*, 27 were *B. burgdorferi* sensu stricto, 30 were *B. garinii*, four were *B. valaisiana*, and two were *B. japonica*. Deqin county had a distribution of four *Borrelia* spp. except *B. japonica* which was only found in Yunlong county (Table 1). Additionally, four of five *Borrelia* spp. were detected in *Apodemus draco* (Table 2). The nucleotide sequences of the *B. afzelii* sequences were closely related to the sequence from a patient in China (JX888444.1). All *B. burgdorferi* sensu stricto sequence were identical to the sequence from strain BRE-13 sequenced from a patient’s CSF in France (KY594010.1). *B. garinii* sequences in this study showed 99% identity with the strain YN12/2012 from *Canis familiaris* in Yunnan Province. *Borrelia japonica* sequences showed 99% identity with strain Cow611C from a tick in Japan (L30125.1). The *B. valaisiana* sequences were similar to the strain KM2 from *Ixodes granulatus* ticks in Taiwan, China (98%, HM100110.1) and the strain CKA2a from *Apodemus agrarius* in Zhejiang,China (99%, AB022124.1). Phylogenetic analyses based on different representative sequences in this study revealed that all detected *Borrelia* fell within five separate clades belonging to five different types of BBSL including *B. afzelii, B. burgdorferi* sensu stricto, *B. garinii, B. japonica* and *B. valaisiana* (Fig 1).

**Fig 1.**
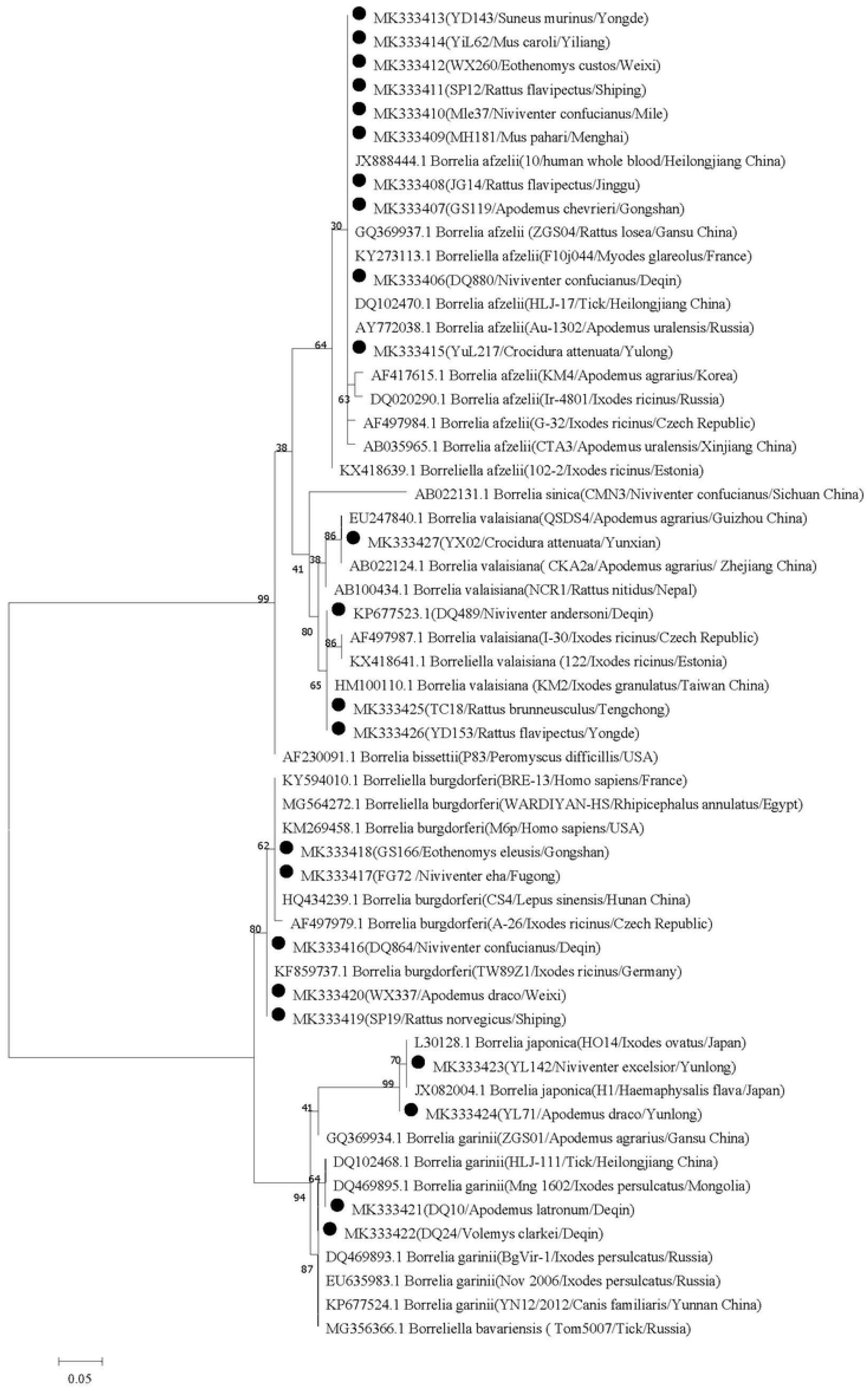
Maximun Likelihood phylogenetic tree based on a comparison of *Borrelia burgdorferi* sensu lato 5S-23S rRNA intergenic spacer gene sequences obtained from Yunnan small mammals with *Borrelia burgdorferi* sensu lato reference strains. The number on each branch shows the percent occurrence in 1000 bootstrap replicates. Black circles stood for novel sequences identified in this study.

## Discussion

Human cases of LB have been confirmed in almost every province found on mainland China including Yunnan Province. However, most of patients only had serological evidence and were not confirmed for specific genotypes. BBSL has been reported in small mammals trapped in the provinces Qinghai, Hunan, Shanxi, Liaoning, Sichuan, Fujian, Zhejiang, Gansu, Guangdong, Jilin and Yunnan [8–16], suggesting that small mammals are likely the main reservoir hosts in China. This study presents a large sample size extending over a wide geographic area, which provides insight into the prevalence, spatial distribution and genetic diversity of BBSL in small mammals collected in Yunnan Province.

We documented BBSL infection in 30 species of small mammal, among which, 20 species had not been previously documented. These species may be infected occasionally, whether they serve as reservoir hosts need a further study. The *Rattus tanezumi* (573/3659,15.66%) was the predominant species trapped in households in Yunnan. *Apodemus draco* (579/3659,15.82%) and *A. chevrieri*(402/3659,10.99%) were the predominant hosts species in Yunnan, which was consistent with results from Europe where *Apodemus* are considered a major reservoir of *Borrelia* [17]. BBSL was detected in *Apodemus draco* and in *A. chevrieri* in Yunnan, with *A. draco* capable of carrying four *Borrelia* spp. The *Ochotona gloveri, Soriculus leucops* and *Rattus tuekkestanicus* also had a much higher prevalence(>14%) with much larger sample sizes in this study than in other provinces in China [12,18–22]. *Rattus norvegicus* is the prominent household species in Yunnan, which had a high prevalence(12.50%) and was detected positive for pathogenic genotypes (*B. afzelii* and *B. burgdorferi* sensu stricto). We also found that the uncommon species *Sorex cylindricauda* in this study tested positive for BBSL DNA, requiring further investigation to fully understand their role in maintaining or amplifying infections in nature.

Our findings indicated that prevalence rates in rodents are ranked highest to lowest by landscape type as follows: forest landscape > agricultural landscape > residential landscape, which is likely related to tick vector density and preferred habitat. This reiterates the need for individuals traveling into potential tick habitats, like the forest, to take proper protective measures to limit tick bite exposure. Sampling locations in this survey contained a broad range of altitudes from 500 meters to 4500 meters. Among the three altitude classes, small mammals with the highest prevalence of BBSL were found above 2500m. It was reported that *Ixodes ricinus* distribution in Sumava National Park extended toward higher altitudes, probably in relation to warming climates[23]. The roles temperature and humidity play in tick reproduction and reservoir preferences requires further investigation within these altitude ranges. Additionally, there are no reported human cases at these heights, which might reflect lower populations living in these areas.

Our study found five genospecies of BBSL in small mammals in Yunnan Province, four of them except for *B. japonica*, have previously been associated with LB [24–25]. There exists a wide distribution and genetic diversity of BBSL in Yunnan, compared to only 1-2 genospecies of BBSL in most provinces in China, such as Qinghai, Zhejiang, Guizhou and Guangxi. According to the sequence analysis carried out in this study, most of the *B. afzelii* sequences shared 99% identity with clinical isolates from patients in northeastern China [26]. Most of the *B. burgdorferi* sensu stricto sequences were identical to the sequence from a human case reported in France (KY594010.1). At this time, there have been no confirmed patients with registered sequence of Lyme disease spirochetes in Yunnan province, requiring further investigation in the near future. The sequence of *B. valaisiana* obtained from small mammals cluster into two clades, one cluster within the sequence from Guizhou and Zhejiang province, the other three cluster fell within close proximity to sequences from Europe. Birds are major reservoirs for *B. valaisiana* in Europe, however the transmission cycle maintaining *B. valaisiana* in Yunan may be different from other areas, requiring additional study. *B. japonica* have only been found in Yunlong county, with this representing the first report documenting *B. japonica* in *Apodemus draco* and *Niviventer excelsior* in China. *B. garinii* is the most common genospecies in China, followed by *B. afzelii* [27]. However, we found that *B. afzelii* was the main genospecies detected in Yunnan, which is consistent with previous reports [4]. *B. burgdorferi* sensu stricto has been detected in *Sika deer* from Jilin and in *Caprolagu ssinensis* from Hunan, and detected in small mammals in Yunnan within the more populated counties of Gongshan, Deqin, and Weixi (S1, S2, S5) found in northwestern Yunnan. These findings reflect that Yunnan Province is of particular interest given its diverse topographic range and high level of biodiversity in small mammals that are potential reservoirs for BBSL.

In conclusion, Yunnan Province is an important natural foci of BBSL in China, and given the absence of reported human cases within this region, efforts to expand clinical surveillance are needed immediately.

## Acknowledgements

We thank Professor Zheng-da Gong for identification of rodent species.

